# Collaborative Behavior of Urea and KI in Denaturing Protein Native Structure

**DOI:** 10.1101/2019.12.29.888271

**Authors:** Qiang Shao, Jinan Wang, Weiliang Zhu

**Affiliations:** Drug Discovery and Design Center, Shanghai Institute of Materia Medica, Chinese Academy of Sciences, 555 Zuchongzhi Road, Shanghai, 201203, China

## Abstract

In this work, the combined influence of urea and KI on protein native structure is quantitatively investigated through the comparative molecular dynamics simulations on the structural dynamics of a polypeptide of TRPZIP4 in a series of urea/KI mixed solutions (urea concentration: 4M, KI salt concentration: 0M-6M). The observed enhanced denaturing ability of urea/KI mixture can be explained by direct interactions of urea/K^+^/water towards protein (electrostatic and vdW interactions from urea and electrostatic interactions from K^+^ and water) and indirect influence of KI on the strengthened interaction of urea towards protein backbone and side-chain. The latter indirect influence is fulfilled through the weakening of hydrogen bonding network among urea and water by the appearance of K^+^–water and I^—^urea interactions. As a result, the denaturing ability enhancement of urea and KI mixed solution is induced by the collaborative behavior of urea and KI salt.

## Introduction

Proteins are marginally stable and are hence very sensitive to environmental conditions. As a highly concentrated waste product in mammalian kidneys, Urea ((*NH* _2_)_2_ *C* = *O*) exerts significant denaturing influence on the native structures of proteins and interrupt their biological functions. The molecular mechanism underlying urea-induced protein denaturation has been extensively studied in the last several decades. Accordingly, “indirect” and “direct” mechanisms have been proposed. The “direct” mechanism proposes that urea unfolds proteins through its direct interactions, which might include the direct hydrogen bonding from urea to protein backbone,^1-3^ electrostatic interactions with polar residues,^4-6^ and van der Waals attraction with residues at protein surface.^7-9^ The importance of each abovementioned urea-protein interaction has been highlighted in these studies, but which interaction contributes dominantly to urea-induced protein denaturation is not confirmed yet. On the other hand, the “indirect” mechanism suggests that the solvation of urea molecules disrupts water structure and thus affects the behavior of the solvent molecules, thus weakening the hydrophobic effects and making the protein hydrophobic groups more readily solvated.^10-14^ As multiple theoretical and experimental studies seemingly support the former mechanism,^1-3,5-9,13,15-17^ others argue that both mechanisms are involved in urea denaturation.^4,9,18-20^

Besides urea denaturation, the studies on the combined effects of urea and other chemical agents, e.g., other natural osmolytes, on protein structure have been initiated by the observation that mixtures of osmolytes are present in the cell,^21-24^ which raises the question whether their actions on protein stability are independent or synergistic.^25^ So far the mixtures of multiple representative protecting osmolytes with urea have been investigated which indicated the counteraction of denaturing effect of urea by protecting osmolytes including trimethylamine N-oxide (TMAO), betaine, sarcosine, and trehalose.^22,26-32^ In addition, to figure out the interplay of two denaturing osmolytes in the mixed solution and their combined effects on protein native structure, Zhou and co-workers studied the structure stability of two proteins (hen egg-white lysozyme and protein L) in the mixture of urea and guanidinium chloride (GdmCl) using molecular dynamics (MD) simulation.^33^ The two denaturants were found to compete with each other to interact with protein and GdmCl has a greater tendency to accumulate on protein surface than urea does. The accumulation of two denaturants on protein surface creates a more crowded local environment around protein and results in unexpected structure collapse of denatured protein, different to the extended unfolded structure observed in isolated GdmCl solution and urea solution.

The influence of inorganic salts on biological function is ubiquitous and specific ion effects can be very important in living organisms.^34^ As organized in Hofmeister series, the species on the left are referred to as kosmotropes which tend to precipitate proteins, while those on the right are called chaotropes which enhance protein solubility and favor protein denaturation. With the knowledge that both urea and chaotropic salt can destabilize and denature proteins, it is interesting to speculate whether the mixed urea and a representative chaotropic salt (e.g., potassium iodide (KI)) has the enhanced denaturing ability compared to the solution containing either similarly concentrated single chemical denaturant. To answer this question, we ran comparative MD simulations previously to measure the structure stability of a β-structured polypeptide (TRPZIP4) in urea/KI mixed solution as well as in similarly concentrated single-component urea solution and KI solution, respectively.^35^ The β-hairpin structure can be partially unfolded in the mixed solution containing both 4M urea and 3M KI whereas the same structure displays strong stability in single-component urea or KI solution, suggesting that the mixture of urea and KI has enhanced ability to denature protein.

In the present study, we ran a series of MD simulations for TRPZIP4 solvated in the solutions containing constant concentration of urea (4M) but varied concentration of KI salt (0M-6M) to further investigate the combined effects of urea and KI on protein structure. In addition, to demonstrate the validity of MD simulation in describing the behavior of urea and KI, two different poplar molecular force fields of urea, namely OPLS^13^ and KBFF^36^ force fields, were utilized in the present study. These comparative studies provide comprehensive, atomic-level picture of the collaborative behavior of urea and KI in denaturing protein native structure.

## Materials and Methods

All-atom MD simulations were performed in explicit solvent at room temperature, making use of AMBER 9.0 suite of programs.^37^ Protein atoms were modeled by AMBER FF99 force field^38^ and water was described with the SPC/E model.^39^ The parameters of the force fields concerning urea were taken from Duffy *et al*. (OPLS force field).^13^ In addition, the KBFF force field of urea developed by Weerasinghe and Smith^36^ was also applied and the corresponding simulations were run as control tests. The parameters for K^+^ and I^-^ ions were taken from those of the non-polarizable spherical ions created by Joung and Cheatham (*ions08*.*lib* in AMBER).^40,41^

In each simulation system of urea/KI mixed solution, TRPZIP4 polypeptide was solvated into a cubic box containing a large amount of solvent (water) and co-solvent (urea, K^+^ and I^-^ ions). The detailed numbers of solvent and co-solvent molecules are listed in Table 1. For each system, the simulation procedure included the energy minimization, the following heating-up process, and the final longtime equilibrium simulation calculation (production). NPT (number, pressure, temperature) ensemble calculations were performed and periodic boundary conditions were used in the simulations. The SHAKE algorithm^42^ was used to constrain all bonds involving hydrogen atoms. A cutoff of 10.0 Å was applied for nonbonding interactions. The Particle Mesh Ewald method was applied to treat long-range electrostatic interactions.^43^

**Table 1.** Parameters for individual simulations of TRPZIP4 polypeptide in urea/KI mixed solutions with varied concentrations of KI. ^a^The parameters in brackets are for the simulations using KBFF force field of urea.

| KI<br>Concentration | Urea | K<sup>+</sup> | I<sup>-</sup> | Water | Simulation<br>Length (ns) |
| --- | --- | --- | --- | --- | --- |
| 0 M | 877 | 0 | 0 | 7238 | 225 |
| 1 M | 877 | 200 | 200 | 7515 (7551)<sup>a</sup> | 224 (248) |
| 2 M | 877 | 400 | 400 | 7220 (7247) | 242 (284) |
| 3 M | 877 | 600 | 600 | 7333 (7320) | 250 (247) |
| 4 M | 877 | 800 | 800 | 7679 (7666) | 235 (286) |
| 5 M | 877 | 1000 | 1000 | 7679 (7666) | 220 (240) |
| 6 M | 877 | 1200 | 1200 | 7679 (7664) | 232 (258) |

The energy of each system was minimized through a total of 2500 steps of calculations: 1000 steps of steepest descent minimization with the polypeptide being fixed using harmonic restraints (using a force constant of 10.0 kcal mol^-1^ Å^-2^ to apply to the backbone atoms), which was then followed by 1500 steps of conjugate gradient minimization. Subsequently, to better relax the system, the system was heated to 360 K and equilibrated for 2 ns at 360 K, followed by a 2 ns of cooling from 360 to 300 K performed with harmonic restraints (force constant = 10.0 kcal mol^-1^ Å^-2^) applied to the backbone atoms. Finally, longtime production runs were performed with the step-size of 0.002 ps, and the data were collected every 2.0 ps.

## Results

### Enhanced Protein Denaturing Ability of Urea Solution in the Presence of KI Salt

Our previous studies showed that during certain simulation time (∼250 ns), the native structure of TRPZIP4 keeps stable in either 4 M urea solution or 3 M KI solution but is largely denatured in the mixture containing both 4M urea and 3M KI. To confirm the enhanced protein denaturing ability of urea in the presence of KI salt, in the present study, we measured the structure stability of the same polypeptide in a series of urea/KI mixed solution with constant concentration of urea (4M) but varied concentration of KI. The time series of root-mean-square deviations (RMSDs) of TRPZIP4 with respect to its native structure in individual solutions are depicted in Figure 1. One can see that the RMSD value keeps steady in low range as the concentration of KI is low (e.g., ≤1M). While the KI concentration is increased, the RMSD value is enlarged, indicating the destabilization of protein structure in high-concentrated urea/KI mixed solution. The final structure of the polypeptide in high-concentrated urea/KI solution is shown in Fig. 2. In comparison to the stable β-hairpin structure in pure water, single-component urea or KI solution, the structure in high-concentrated urea/KI is largely unfolded, with the breaking of most of backbone native hydrogen bonds.

**Figure 1.**
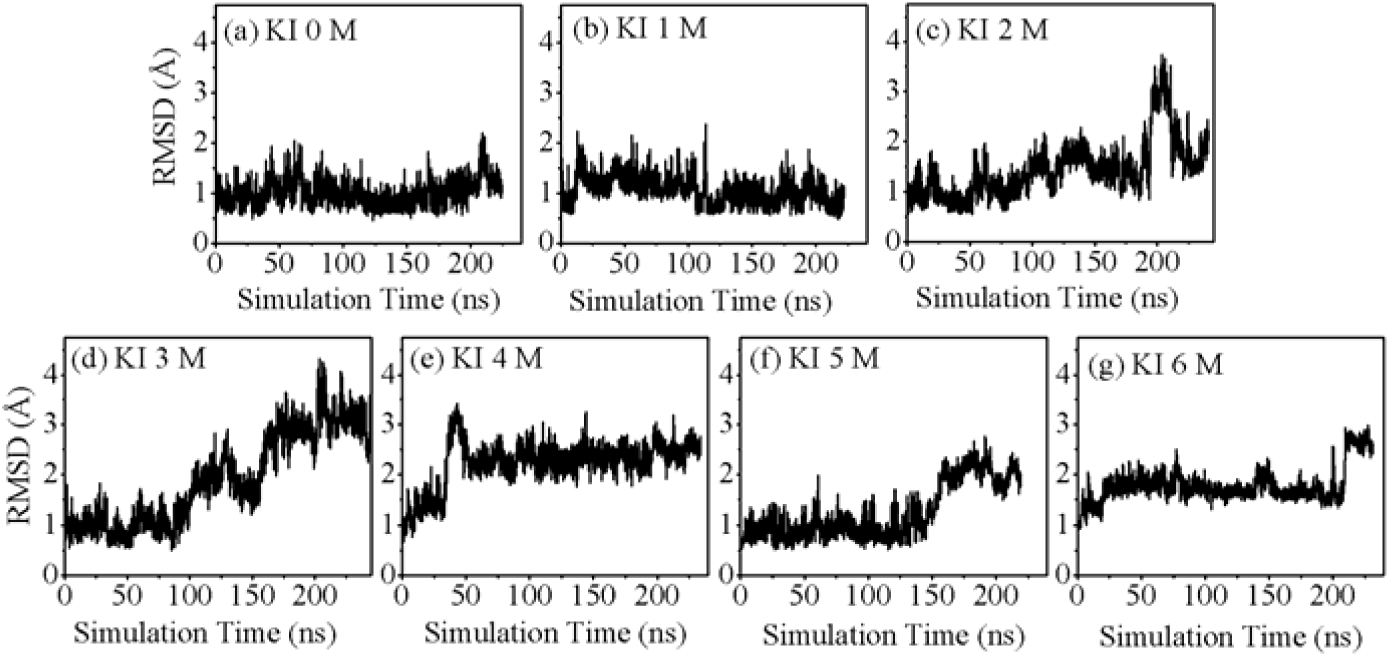
The root-mean-square deviation (RMSD) of TRPZIP4 polypeptide corresponding to its NMR structure as a function of simulation time in a series of urea/KI mixed solutions with varied concentrations of KI.

**Figure 2.**
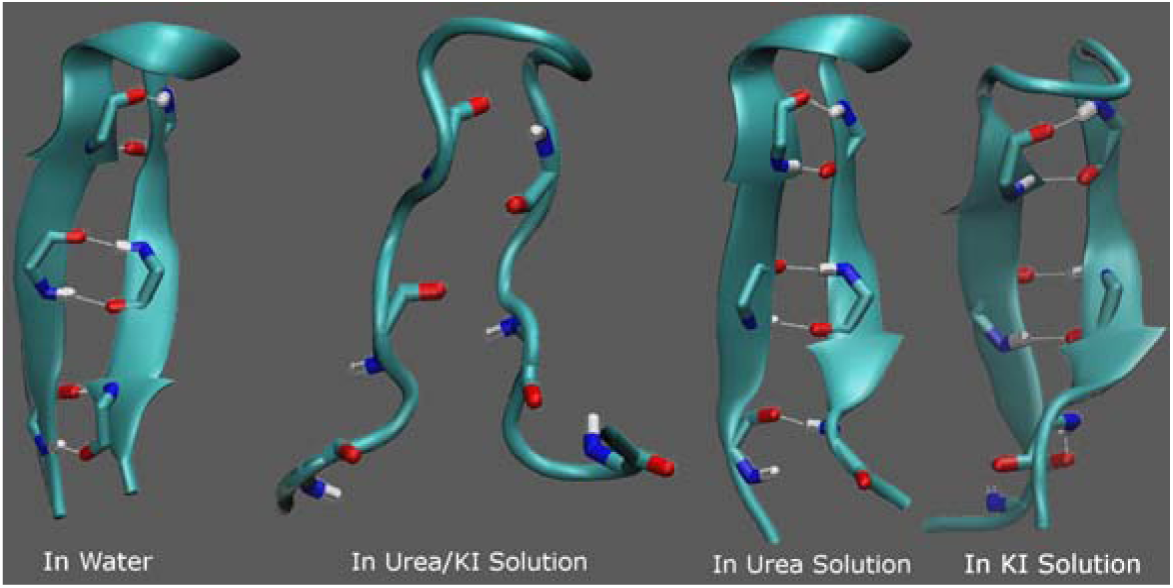
Simulated structures of TRPZIP4 in high-concentrated urea/KI mixed solution and the comparison to the corresponding stable structure in pure water, urea, or KI solution. The backbone native hydrogen bonds are represented by dash lines.

### Microscopic Solvent Environment around Protein Surface

Many previous studies showed that urea has a strong tendency to approach protein.^18,20,44,45^ To see the microscopic solvent environment around protein, we calculated the average number of solvent and co-solvent molecules in the first solvation shell (FSS) of protein (the area of 3.4 Å around protein surface) in all urea/KI mixed solutions under study. In addition, the molar ratio of urea and water molecules around the protein surface relative to the urea/water ratio in the entire simulation system is used to evaluate the preferred binding of urea to protein surface 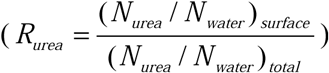: urea accumulates around protein if the relative ratio is greater than 1; otherwise, it is disposed from protein surface.

One can see from Figure 3 that the number of urea molecules staying in the FSS of protein only slightly fluctuates, the numbers of K^+^ and I^-^ ions are increased gradually, but the number of water is apparently decreased with the increasing of KI concentration. The relative ratio *R*_*urea*_ of urea is significantly larger than 1 even in the absence of KI, indicating that urea molecules heavily accumulate around protein. As the KI concentration is increased in mixed solution, more K^+^ and I^-^ ions approach protein surface and thus removes some of the surrounding water molecules. As a result, although the number of urea molecules in the FSS of protein is little changed, the value of *R*_*urea*_ keeps increasing with the addition of KI (Figure 3 (a)), showing the increasing of urea accumulation level as the KI concentration is increased. On the other hand, K^+^ cations also have a tendency to accumulate around protein whereas I^-^ more likes to be repelled from protein and solvated in bulk solution (Figure 3 (c) and (d)).

**Figure 3.**
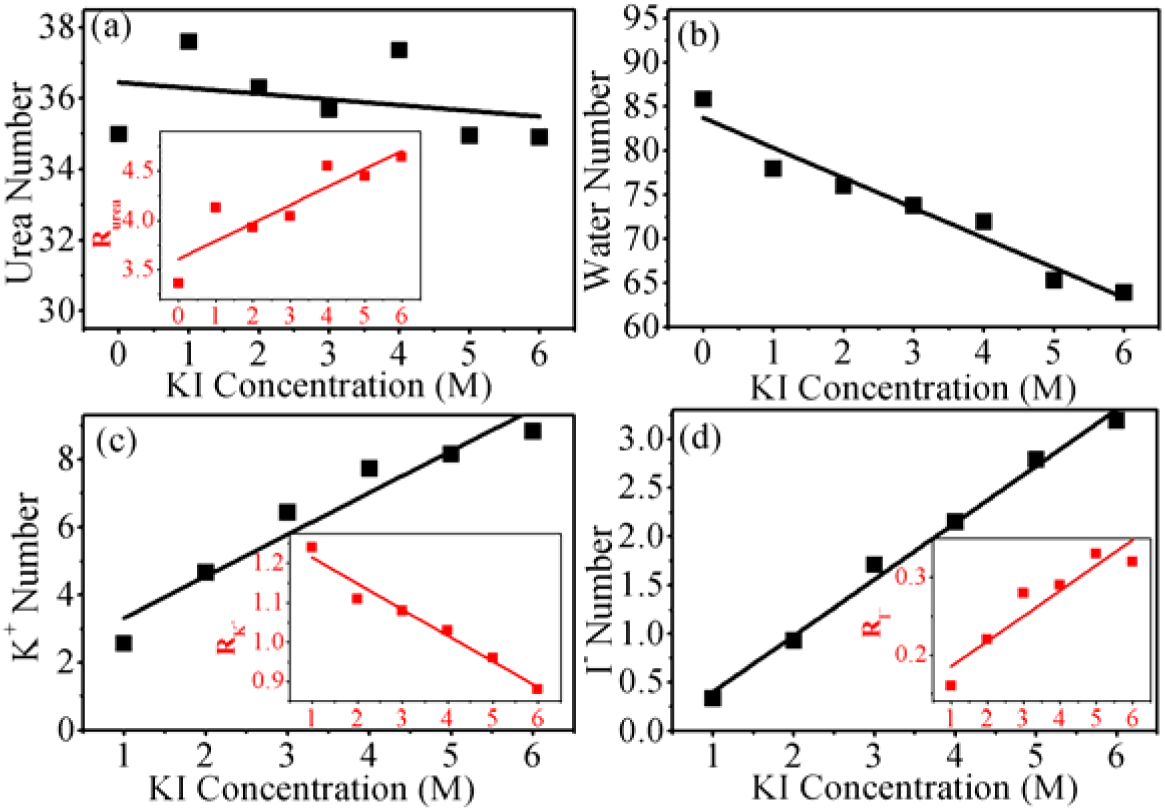
KI concentration dependence of the average number of (a) urea, (b) water, (c) K^+^, and (d) I^-^ in the first solvation shell of TRPZIP4 polypeptide in urea/KI solutions. The insets show the corresponding relative ratio of urea, K^+^, and I^-^.

### Effects of KI Concentration on the Molecular Interaction of Solvent to Protein

As the donor and acceptor of backbone hydrogen binding, the backbone carbonyl (-C=O) and amide (-NH) groups are subject to the external interactions from solvent molecules, which eventually influences the stability of protein backbone hydrogen bonds. The damage of the TRPZIP4 structure in urea/KI solution is mainly the backbone hydrogen bond breaking. To understand the mechanism of urea and KI induced protein denaturation, we first analyzed the molecular interactions between protein backbone and water/urea/K^+^ (I^-^ anions are rather expelled from protein, whose interaction is thus not investigated hereinafter). Figure 4 shows the radial distribution function (RDF) of the oxygen and nitrogen atoms of urea, the water oxygen, K^+^ cation around protein backbone carbonyl and amide groups in various urea/KI solutions. The data were collected from the last 50 ns of the simulations in the respective solutions.

**Figure 4.**
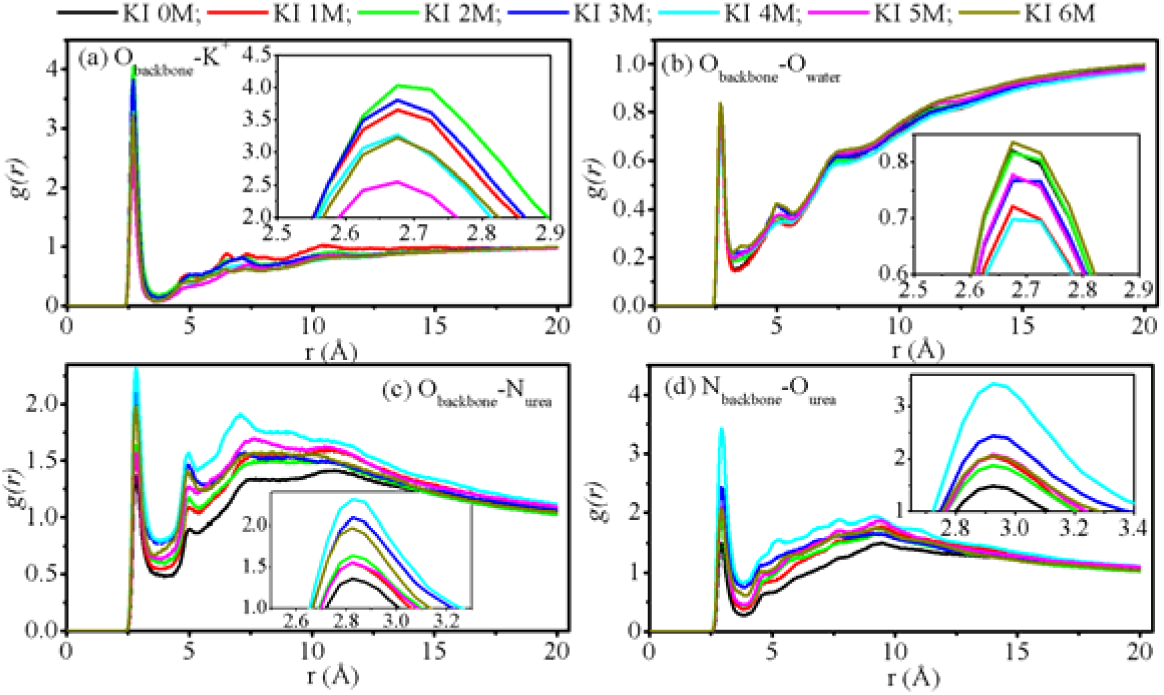
RDF for K^+^ cation, oxygen atom of water, oxygen and nitrogen atoms of urea around backbone carbonyl and amide groups in various urea/KI solutions. O_backbone:_ backbone carbonyl O oxygen; N_backbone_: backbone amide nitrogen atom; O_water_: water oxygen atom; N_urea_: urea nitrogen atom; O_urea_: urea oxygen atom.

A high peak can be seen in the RDF diagram of K^+^ around protein backbone carbonyl group (at *r* ≈2.68 Å in Figure 4 (a)). Meanwhile, the peak of K^+^ around protein backbone amide group is very low (data is not shown here). Therefore, K^+^ could have a strong electrostatic interaction with backbone carbonyl but not amide group. The strong binding affinity of K^+^ cation (but not I^-^ anion) to protein backbone has been observed previously. Several MD simulations observed that while alkali cations (Na^+^, K^+^ et al.) exhibit affinity to backbone carbonyl and side chain carboxylate oxygens, none of the counterions (halide anions) shows appreciable affinity to protein amide hydrogen.^46-49^ Therefore, the binding of cation to backbone has been considered by these authors as the main reason for chaotropic salt induced protein denaturation.^46-51^ In addition, the strong binding affinity of a series of metal cations to carboxylate group was demonstrated in the LCST measurement and vibrational sum frequency spectroscopy (VSFS) experiment.^52^

High peaks can be also seen in the RDF diagrams of the water oxygen and urea amide nitrogen around protein backbone carbonyl group (at *r*≈2.8 Å in Figure 4 (b and c)), demonstrating the formation of hydrogen bonds from the hydrogen bond donor of water and urea to the acceptor of backbone carbonyl group. In addition, there is also a peak for the urea carbonyl oxygen around backbone amide group (at *r*≈3.0 Å in Figure 4 (d)), which implies that urea can also work as a hydrogen bond acceptor and form hydrogen bond with backbone amide group. Similar conclusion was also drawn in multiple previous MD simulation studies. For instance, Gao and co-workers suggested that urea could form hydrogen bonds with carbonyl groups of protein in the denaturation process and, more importantly, form hydrogen bonds with amide groups in the denatured protein structure and therefore stabilizes the denatured state.^18-20^

Different influence of KI concentration can be seen for the peak heights representing different protein-solvent interactions. As shown in Figure 4, the peak height representing K^+^-backbone carbonyl group interaction is lowered as the KI concentration becomes larger (e.g., >3 M). Meanwhile, although the peak height representing the interaction of water oxygen and backbone carbonyl group changes little, the peak height for the RDF of either urea nitrogen to protein carbonyl group or urea oxygen to protein amide group roughly follows the order from low to high as the KI concentration is increased, consistent with the enhanced denaturing ability of high-concentrated urea/KI solution.

The RDF diagrams of the oxygen and nitrogen atoms of urea, the water oxygen, and K^+^ cation around protein side-chain (SC) in all urea/KI solutions are also depicted. Figure 5 (a) demonstrates the presence of strong interactions between K^+^ and protein side-chain (mainly the electrostatic interactions between K^+^ and negatively charged side-chain). The peak height corresponding to the interaction between K^+^ and protein side-chain is slightly lowered as the KI concentration is increased, following the same order of the RDF peak height representing K^+^-backbone carbonyl group interaction. The RDF peak of water oxygen around protein side-chain is low, indicating the weak interaction between the two species (Figure 5 (b)). Urea, on the other hand, has strong interaction with protein side-chain ((Figure 5 (c) and (d)), with the corresponding peak heights increased with KI concentration.

**Figure 5.**
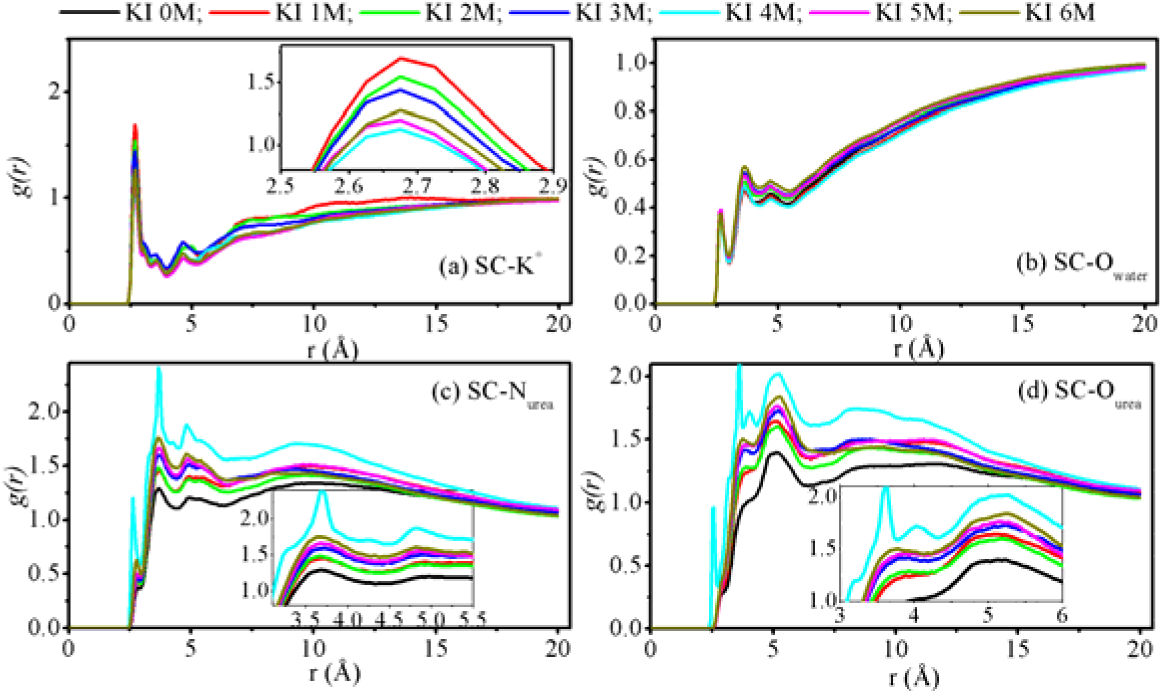
RDF for K^+^ cation, oxygen atom of water, oxygen and nitrogen atoms of urea around side-chain (SC) of TRPZIP4 peptide in various urea/KI solutions.

The preferential interactions between protein and urea, water, and K^+^ can be explained from energetic perspective. The electrostatic energy and van der Waals (VDW) interaction energy between each urea/K^+^/water in proximity to protein (defined as the area within 5.0 Å of protein) and in bulk region (6.0 Å away from any protein atoms), respectively, with the rest of the system were calculated.^8,53^ A spherical cutoff of 13.0 Å was applied in VDW potential energy calculation and no cutoff was applied for long-range electrostatic interactions. As shown in Figure 6, the electrostatic energies of all single solvent (water) and co-solvent (urea and K^+^) molecules are negative. In addition, the electrostatic energies in the region of protein surface are more negative than the counterparts in bulk region. Therefore, urea, water, and K^+^ could have favorable electrostatic interactions with protein. On the other hand, the VDW energy of single urea is also negative, and the VDW energy around protein is even more negative than that in bulk region, suggesting that urea could also has favorable VDW interactions with protein. In contrast, the VDW energy of either water or K^+^ is positive, reflecting the unfavorable VDW interactions of the two species. The increasing of KI concentration in solution does not influence the electrostatic energy of single K^+^ in bulk solution but does increase the electrostatic energy of single K^+^ around protein (Figure 6 (c)). In addition, the increasing of KI concentration also influences the electrostatic and VDW energies of single urea and water in the regions of protein surface and bulk solution: as the KI concentration is increased, the electrostatic energy of single urea is reduced but the electrostatic energy of single water is increased slightly; meanwhile, the VDW energy of single urea is slightly increased.

**Figure 6.**
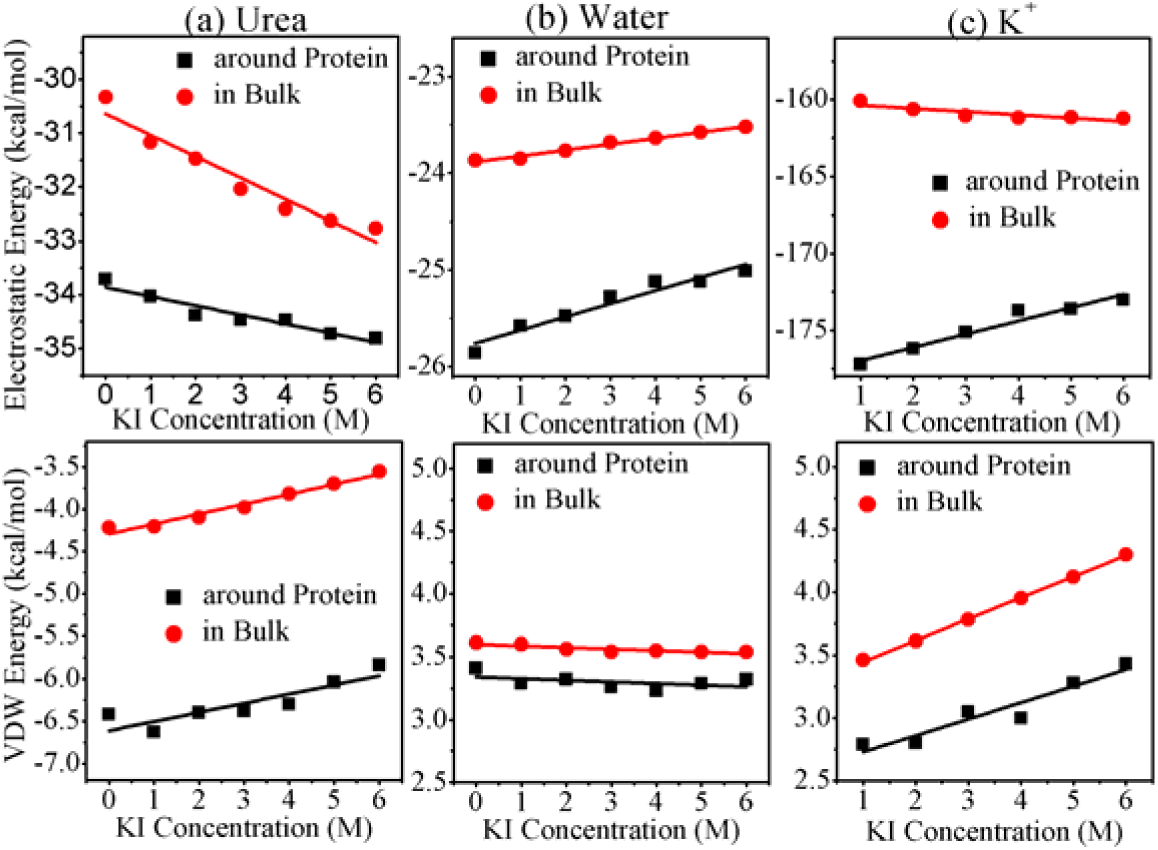
KI concentration dependence of (a-c) electrostatic energy and (d–f) van der Waals energy of each urea/K+/water in proximity to protein and in the bulk region, respectively, with the rest of system for TRPZIP4 in urea/KI mixed solutions.

Figure 7 shows the decomposition of electrostatic and VDW energies per urea/water/K^+^ to protein into protein backbone and side chain parts. As shown in this figure, the electrostatic interactions of all solvent and co-solvent molecules with protein are more biased to protein side-chain rather than backbone, as revealed by more negative electrostatic energy with protein side-chain. As the KI concentration is increased, the electrostatic energy of either single urea or single water molecule to protein is increased and the energy increasing is mainly on protein side-chain. Meanwhile, the electrostatic energy of single K^+^ cation to protein (either backbone or side-chain) has no clear tendency to change. On the other hand, while urea has also favorable VDW interaction to protein, its VDW interaction is more favorable with protein side-chain. The VDW interaction energy between single urea and protein backbone or side-chain is reduced with the increasing of KI concentration. In contrast, the VDW energies per water or K^+^ to protein is close to 0, again showing the unfavorable VDW interaction between water/K^+^ to protein.

**Figure 7.**
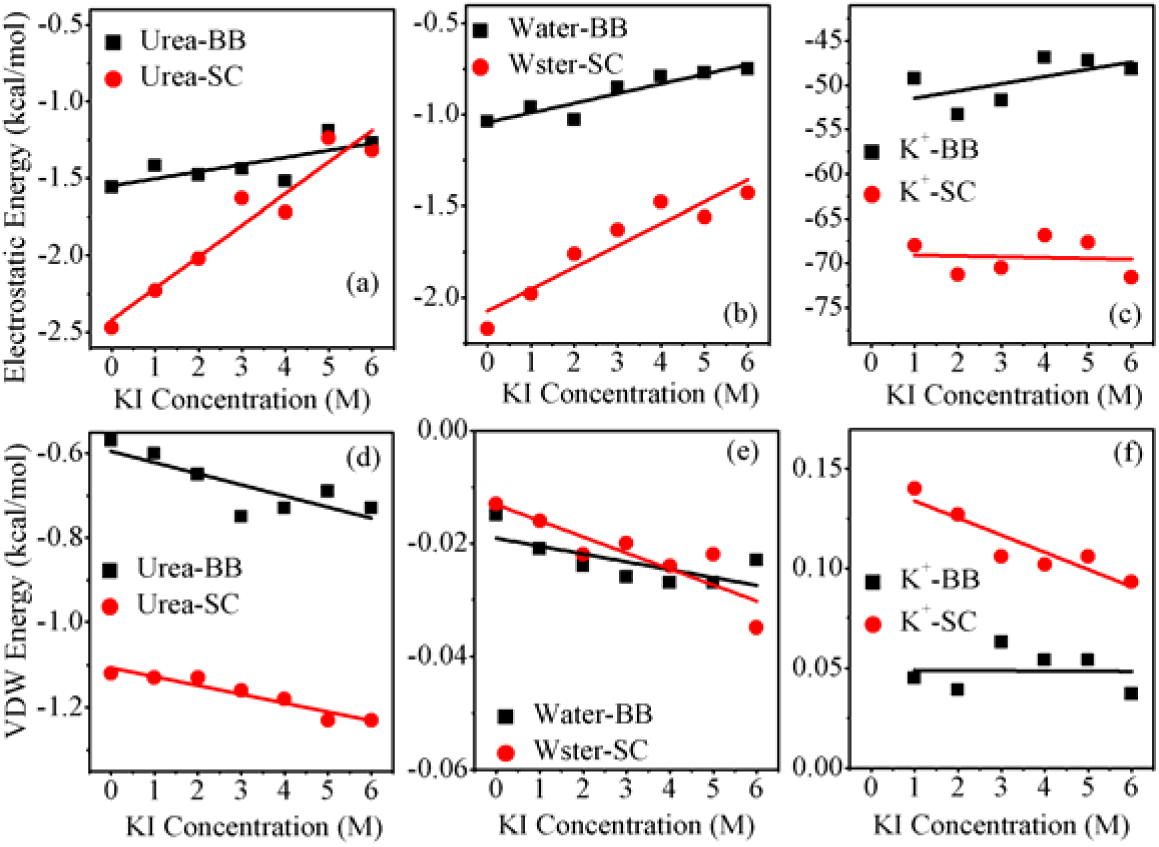
KI concentration dependence of (a-c) electrostatic energy and (d–f) van der Waals energy of each urea/K+/water to protein backbone (BB) or side-chain (SC) for TRPZIP4 in urea/KI mixed solutions.

Molecular force field is crucial for the accuracy of MD simulation. To see the force field influence on simulation results, another popular force field of urea (the KBFF force field)^36^ was also used to run parallel MD simulations for all concentrations of urea/KI mixed solutions. Consistent simulation results were obtained for the two force fields used, showing the convergence of the present simulation study. For instance, as shown in Figure S1 in supporting information (SI), although the exact numbers of urea, water, K^+^ and I^-^ in the FSS of protein are different, the heavy accumulation level of urea and its increasing tendency with KI concentration, the increasing of K^+^ and I^-^ numbers and decreasing of water number in the FSS of protein with the increasing of KI concentration, the moderate accumulation level of K^+^ and the expelling of I^-^ from protein surface can be seen in the simulations of both force fields. In addition, the electrostatic and van der Waals energies of each urea/K+/water with the rest of system in the region of either protein surface or bulk solution (Figure S2), the electrostatic and VDW energies per urea/water/K^+^ to protein backbone and side chain (Figure S3) which were measured from the simulation of KBFF force field change in the same tendency as that from the simulation of OPLS force field.

### Effects of KI Concentration on the Molecular Interactions among Solvent Molecules

While K^+^, urea, and water have favorable interactions with protein, they can interact with each other in bulk solution as well. In Figure 8, we evaluated the K^+^–urea, K^+^–water, K^+^–I^-^, I^—^urea, and I^-^–water distributions in urea/KI solutions. One can see from Figure 8 (a) that K^+^ has a very strong interaction with urea molecule in urea/KI solution, consistent with the observation in previous MD study that urea preferentially solvates positively charged ion.^54^ In addition, K^+^ also has strong interaction with water molecule and I^-^ anion (Figure 8 (b and c)). The peak heights representing K^+^–water and K^+^–I^-^ interactions are increased whereas the peak height representing K^+^–urea interaction is decreased with the increasing of KI concentration, suggesting that the interaction of K^+^ and urea is more likely to be replaced by water solvation or the formation of K^+^–I^-^ ion-pairing in bulk solution in the presence of high-concentration of KI. On the other hand, while I^-^ anion is heavily solvated by water, it can also form strong interaction with urea (see the high peak in Figure 8 (d)) and such interaction is strengthened at high concentration of KI. Finally, urea and water can form hydrogen bonds with each other, and the peak height representing the hydrogen bonding between urea carbonyl oxygen and water hydrogen is decreased whereas the peak height for the hydrogen bonding interaction between urea amide hydrogen and water oxygen is increased with the increasing of KI concentration in urea/KI mixed solution (Figure 9).

**Figure 8.**
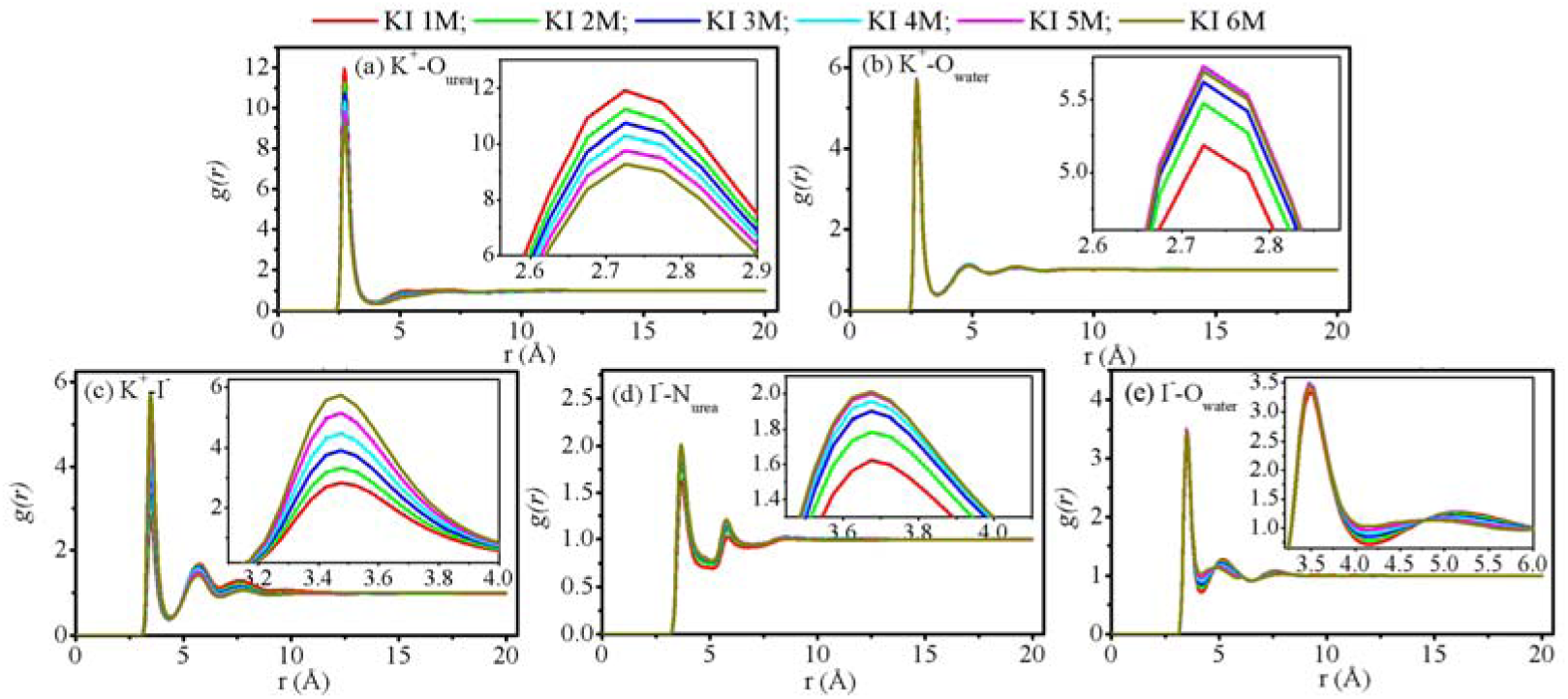
RDF for K^+^ and I^-^ ions around urea and water in various urea/KI solutions.

**Figure 9.**
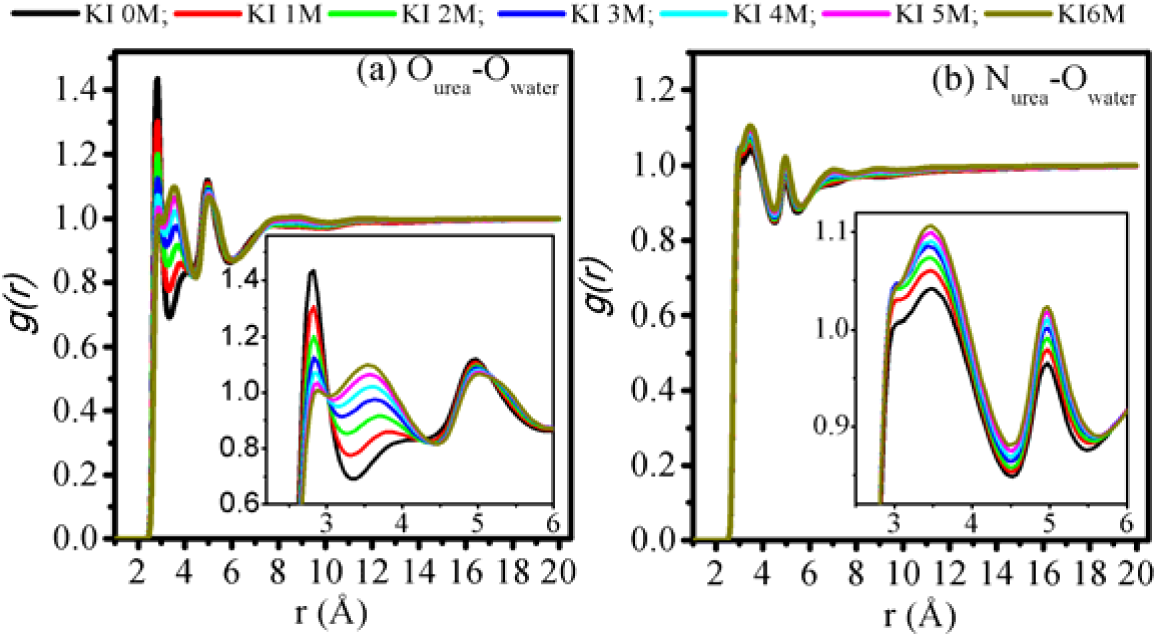
RDF for urea and water around each other in various urea/KI solutions.

## Discussion and Conclusions

In a recent comparative molecular dynamics simulation study, we found that a β-structured peptide has its native structure largely denatured in a mixed solution containing 4M urea and 3M KI whereas the same protein structure keeps significantly stable in similarly concentrated single-component urea solution and KI solution in the same simulation time.^35^ Therefore, it is reasonable to speculate that the presence of KI salt might improve the denaturing ability of urea solution. To figure out the interplay of urea and KI in aqueous solution and their combined effects on protein native structure, in the present study, we performed a series of MD simulations for TRPZIP4 peptide in the solutions containing constant concentration of urea (4M) but varied concentration of KI salt (0M-6M). Two popular OPLS^13^ and KBFF^36^ force fields of urea, were used, which yield similar results, demonstrating the availability of the present MD simulations in describing the behavior of the osmolyte mixed solution.

The detailed analysis of the microscopic solvent environment around protein surface and in the bulk indicates the accumulation of urea and K^+^ cations but not I^-^ anions around protein, creating an environment in which urea could have favorable electrostatic and vdW interactions and K^+^/water also provide favorable electrostatic interactions with protein. The decomposition of these interactions further suggests that these protein-solvent/cosolvent interactions are more biased to protein side-chain rather than backbone. As the intra-protein backbone hydrogen bonds contribute to the stabilization of the global structure of TRPZIP4, urea forms direct hydrogen bonding with protein backbone carbonyl and amide groups, and K^+^ cations and water have strong electrostatic and hydrogen bonding interactions with backbone carbonyl group, respectively, which to some extent impair the backbone hydrogen bonding stability.

In bulk solution, K^+^ could have a very strong interaction with urea molecule, and meanwhile strong interaction with water molecule and I^-^ anion. The RDF analysis shows that K^+^–water and K^+^–I^-^ interaction strength are increased but K^+^–urea interaction is decreased with the increasing of KI concentration, suggesting that the K^+^–urea interaction might be probably replaced by water solvation or the formation of K^+^–I^-^ ion-pairing. Meanwhile, the hydrogen bonding network among urea and water is also weakened as the KI concentration is increased. As a result, the protein-urea interaction is strengthened, which as coupled with the increased protein-K^+^ interactions because of the more replacement of water by K^+^ in the first solvation shell of protein, lead to the enhanced denaturing ability of high-concentrated urea/KI solution. In summary, the enhanced denaturing ability of urea and KI mixed solution is induced by the collaborative behavior of urea and KI salt.

## Acknowledgements

We thank the National Science Foundation of China for finance support (Grants No. 21373258 and 21003003 and 2012AA01A305). The simulations were run at Shanghai Supercomputer Center and TianHe 1 supercomputer in Tianjian.

