## Supplemental Information for "Collaborative Behavior of Urea and KI in Denaturing Protein Native Structure"

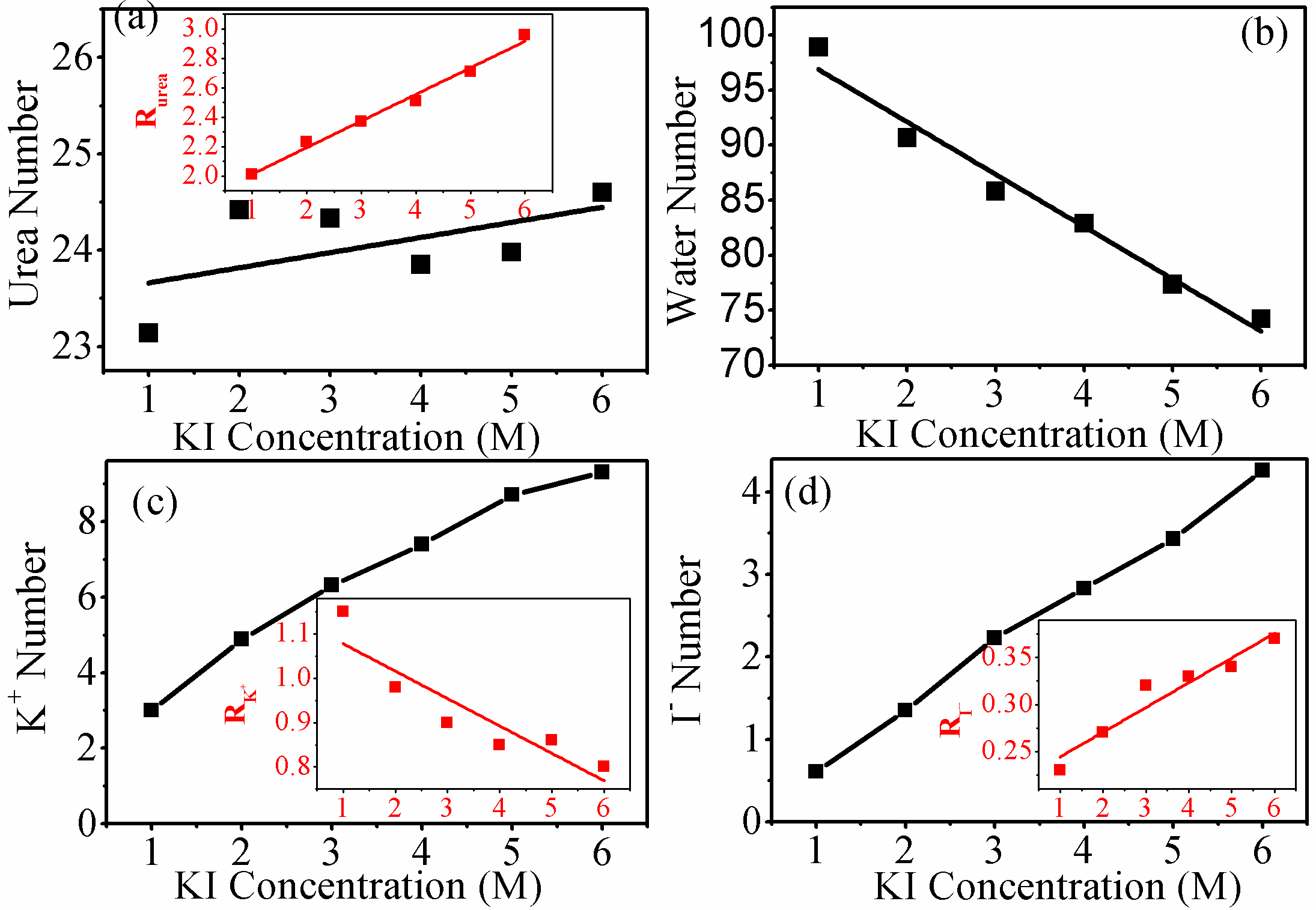

Figure S1. KI concentration dependence of the average number of (a) urea, (b) water, (c) K^+^, and (d) I^-^ in the first solvation shell of TRPZIP4 polypeptide in urea/KI solutions. The insets show the corresponding relative ratio of urea, K^+^, and I^-^. The figures are drawn for the simulations using the KBFF force field of urea.

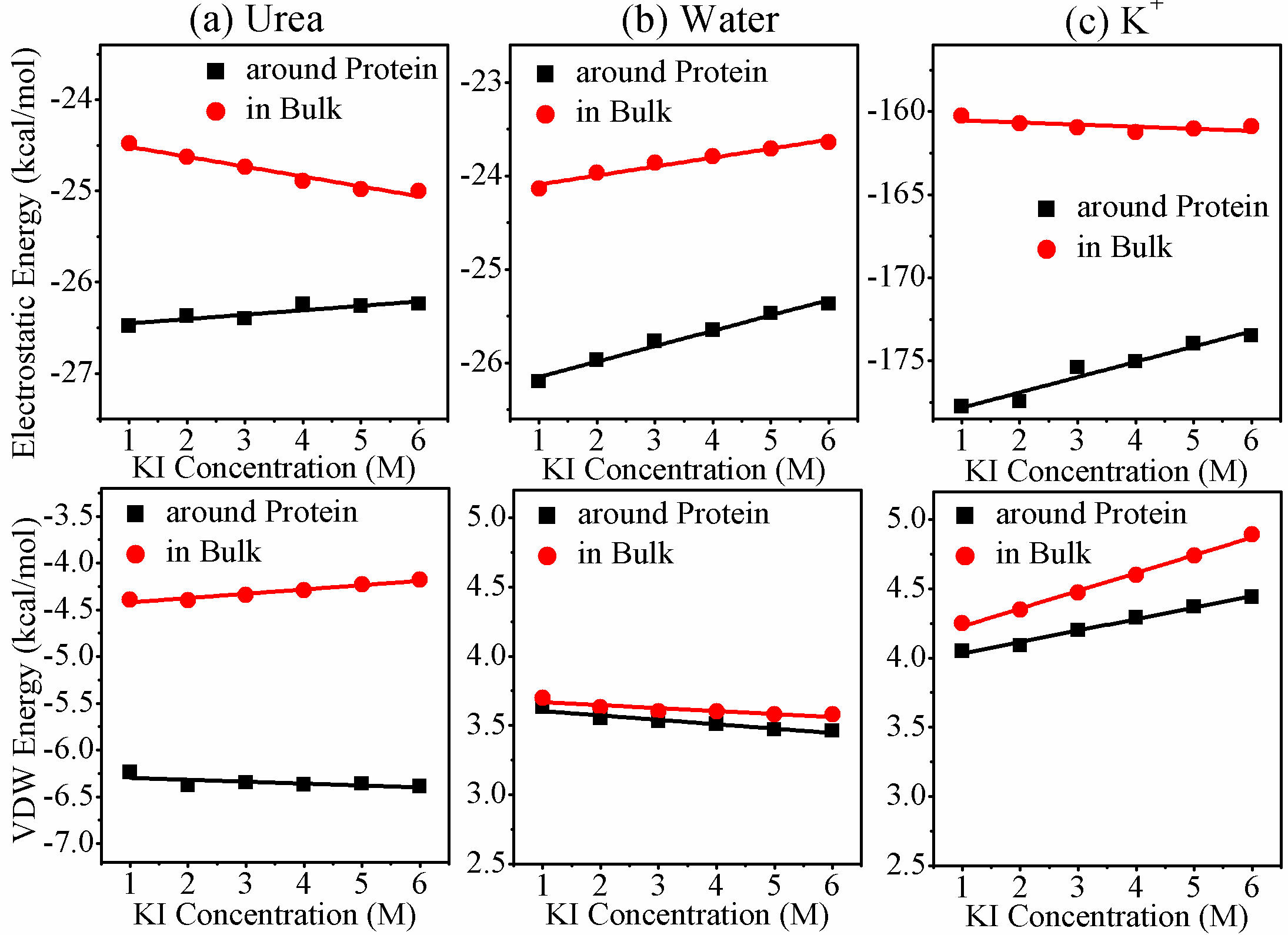

Figure S2. KI concentration dependence of (a-c) electrostatic energy and (d–f) van der Waals energy of each urea/K+/water in proximity to protein and in the bulk region, respectively, with the rest of system for TRPZIP4 in urea/KI mixed solutions. The figures are drawn for the simulations using the KBFF force field of urea.

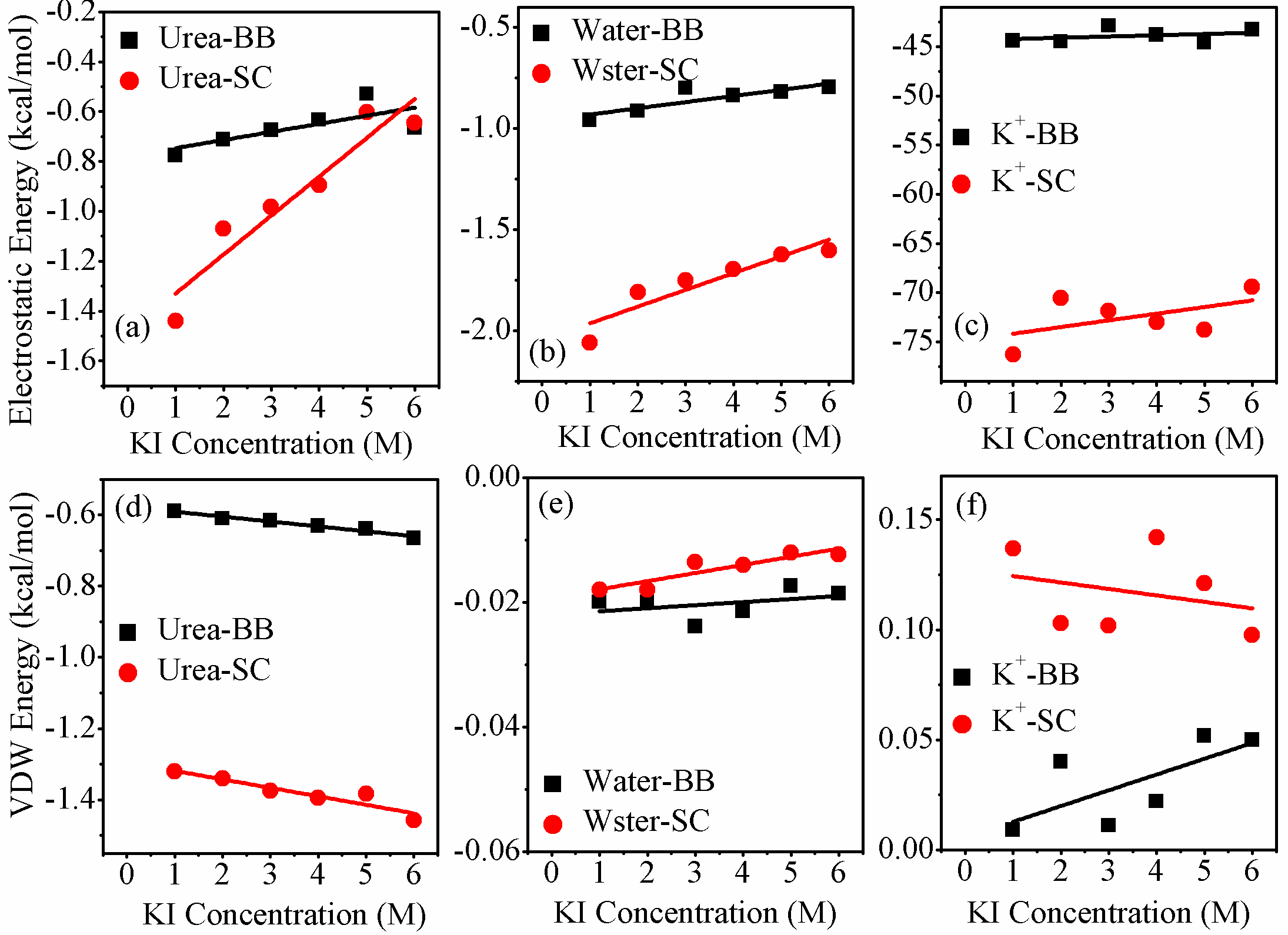

Figure S3. KI concentration dependence of (a-c) electrostatic energy and (d–f) van der Waals energy of each urea/K+/water to protein backbone (BB) or side-chain (SC) for TRPZIP4 in urea/KI mixed solutions. The figures are drawn for the simulations using the KBFF force field of urea.
